# HYPERGLYCEMIA-INDUCED MIR-467 DRIVES TUMOR INFLAMMATION AND GROWTH IN BREAST CANCER

**DOI:** 10.1101/2020.07.01.182766

**Authors:** Jasmine Gajeton, Irene Krukovets, Santoshi Muppala, Dmitriy Verbovetskiy, Jessica Zhang, Olga Stenina-Adognravi

## Abstract

Tumor microenvironment contains the parenchyma, blood vessels, and infiltrating immune cells, including tumor-associated macrophages (TAMs). TAMs affect the developing tumor and drive cancer inflammation.

Hyperglycemic patients have a higher risk of developing breast cancer (BC). We have identified a novel miRNA-dependent pathway activated by hyperglycemia that promotes BC angiogenesis and inflammation supporting BC growth. miR-467 is upregulated in endothelial cells (EC), macrophages, BC cells, and in BC tumors. A target of miR-467, thrombospondin-1 (TSP-1), inhibits angiogenesis and promotes resolution of inflammation. Systemic injections of a miR-467 antagonist in mouse models of hyperglycemia resulted in decreased BC growth (*P*<.001). Tumors from hyperglycemic mice had a 2-fold increase in macrophage accumulation compared to normoglycemic controls (*P*<.001), and TAM infiltration was prevented by the miR-467 antagonist (*P*<.001). BC specimens from hyperglycemic patients had increased miR-467 levels, increased angiogenesis, decreased levels of TSP-1, and increased TAM infiltration in malignant breast tissue in hyperglycemic vs normoglycemic patients (2.17-fold, *P*=.002) and even in normal breast tissue from hyperglycemic patients (2.18-fold inc., *P*=.04). In malignant BC tissue, miR-467 levels were upregulated 258-fold in hyperglycemic patients compared to normoglycemic patients (*P*<.001) and increased 56-fold in adjacent normal tissue (*P*=.008).

Our results suggest that miR-467 accelerates tumor growth by inducing angiogenesis and promoting the recruitment of TAMs to drive hyperglycemia-induced cancer inflammation.

## INTRODUCTION

In the U.S., breast cancer (BC) is the most common cancer diagnosis in post-menopausal women and is the second leading cause of cancer deaths (1). Being overweight or diabetic, combined with low physical activity, are the known risk factors for developing BC in women ages 50 and older (2, 3). There are several meta-analyses studying the association between hyperglycemia, diabetes, and BC, however, the exact mechanisms are not well understood (4–6). Overall, the data suggest that patients with hyperglycemia or pre-existing diabetes have a decreased overall survival and disease-free survival (4).

In 2011, inflammation was added to the “Hallmarks of Cancer” as one of the emerging characteristics of cancer [reviewed by Hanahan and Weinberg (7)]. Chronic inflammation is not only characteristic of obese and T2D patients, but also present specifically in mammary tissue in the majority of obese individuals (8–11). This creates an interesting interaction between macrophages and adipocytes and may be important in understanding BC progression (12, 13). In both adipose and mammary tissues from obese individuals, increased pro-inflammatory gene expression and the characteristic crown-like structures of macrophages were shown in diet-induced and genetic mouse models of obesity (10, 14, 15). Additionally, primary tumors secrete cytokines, such as CCL2, to recruit blood monocytes into the tumor and sustain chronic inflammation.

There, they differentiate into tumor-associated macrophages (TAMs) and shift the milieu to support tumor growth (8, 13). Clinically, the increased infiltration of TAMs has a strong correlation with a poor prognosis (16, 17).

Thrombospondin-1 (TSP-1) is a matricellular protein that inhibits angiogenesis and promotes the resolution of inflammation (18–22). Due to the anti-angiogenic properties of TSP-1, its role in cancer growth has been investigated: TSP-1 has been shown to control tumor growth (23, 24).

We have discovered a novel hyperglycemia-induced pathway that upregulates miR-467 and, as a result, downregulates its target TSP-1 in a cell- and tissue-specific manner, leading to increased angiogenesis (25). Injections of a miR-467 antagonist in mice prevented hyperglycemia-induced BC growth and angiogenesis (26). We have recently found that miR-467 is upregulated by high glucose in cultured macrophages (27). Here, we report the results of a study investigating the role of a hyperglycemia-induced miR-467-dependent pathway in regulation of inflammation in mouse BC and hyperglycemia models and in hyperglycemic patients. miR-467 was dramatically upregulated in BC tissues of hyperglycemic patients and was found circulating in blood. Therefore, we explored the therapeutic potential of this miRNA as a circulating biomarker of BC tumor.

## RESULTS

### Hyperglycemia induces EMT6 cancer growth

In three different mouse models of hyperglycemia, BC growth was accelerated by the high glucose levels (Fig. 1). In a model of Type 1 Diabetes, BALBc mice that were injected with streptozotocin (STZ) to induce hyperglycemia had 2.11-fold increase in tumor weight compared to normoglycemic (citrate buffer injected) controls. (Fig. 1a; 0.585±0.2080 g vs 0.277±0.1213 g, *P*<.001). In *Lepr*^db/db^ mice, a genetically diabetic mouse that models Type 2 Diabetes, tumor weights were increased by 4.22-fold compared to their genetically normoglycemic *Dock7^m^+/Lepr*^db^ heterozygote control group (*Lepr^Dock/db^*) (Fig. 1b; 0.373±0.136g vs 0.088±0.037g, *P*=.0079). In a diet-induced hyperglycemia model, WT C57BL6 mice fed a “Western diet” for 16 weeks was also used to more closely model chronic hyperglycemia in humans. There was a 2.95-fold increase in tumor weights Western diet-fed mice compared to chow-fed controls (Fig. 1c; 0.374±0.157g vs 0.127± 0.047g, *P*<.001) and had significantly elevated glucose levels (129.4±21.44 mg/dL vs 86.35±9.172 mg/dL, *P*<.001).

**Figure 1.**
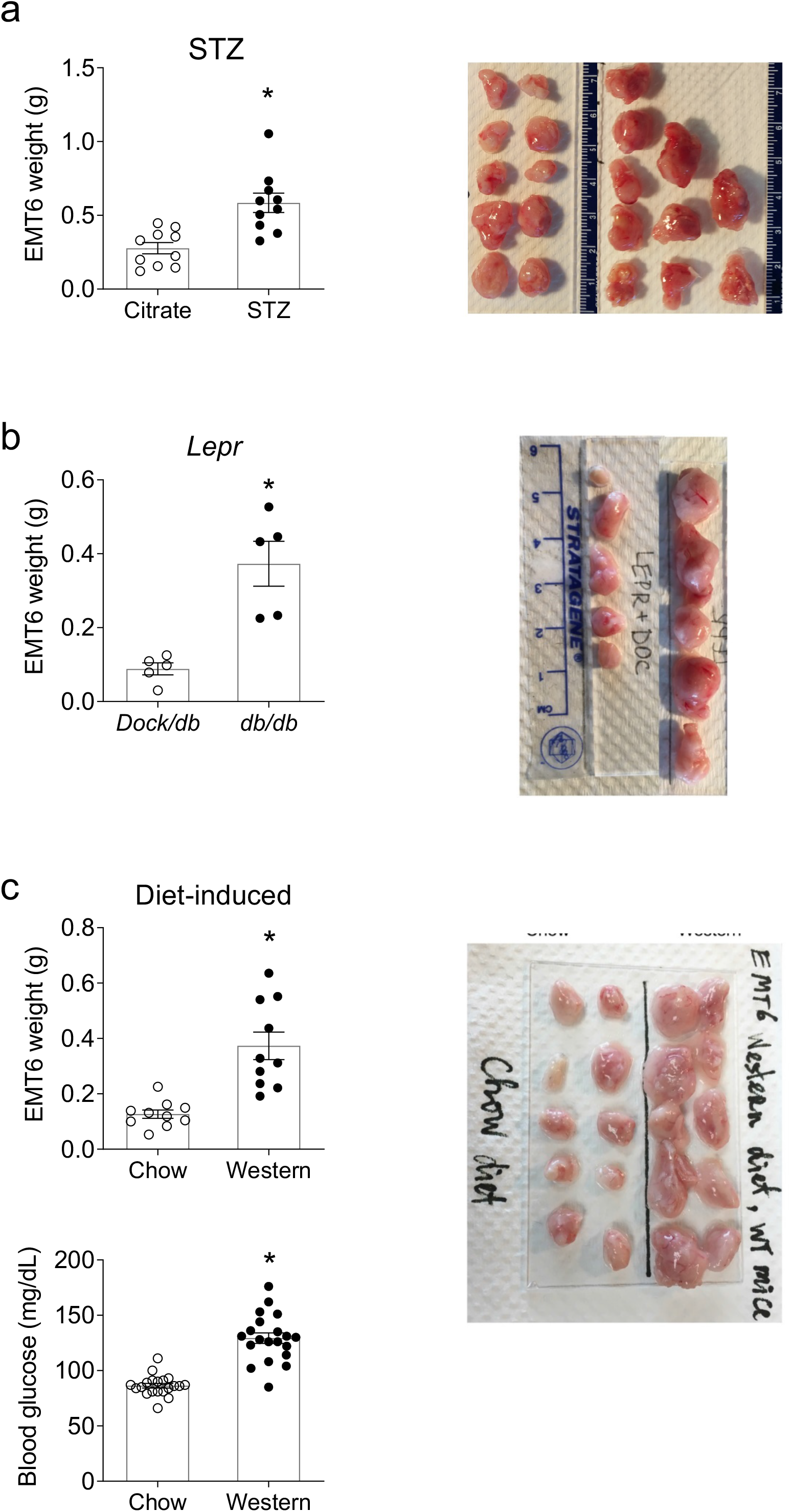
Increased EMT6 BC tumor weights in hyperglycemic mice. EMT6 cancer cells were injected subcutaneously into normoglycemic or hyperglycemic: a. BALB/c mice (n=10/group), b. *Lepr* mice (n=5/group), c. WT mice on long term Western diet (n=10/group), as described in Methods. Fasting blood glucose levels were measured at end point. Tumors were excised a week later and weighed. Bars represent the mean ± SEM. * *P*<.05

### Increased expression of pro-inflammatory markers and markers of macrophages in BC from hyperglycemic mice

Expression of macrophage and pro-inflammatory markers were assayed in EMT6 tumors from hyperglycemic mice (Fig. 2, 3). Expression of pro-inflammatory markers *Il6*, *Ccl2*, and *Tnf* were significantly increased in STZ-treated hyperglycemic mice compared to a normoglycemic, citrate buffer control group (Fig. 2a). To further understand the type of macrophage populations present in these tumors, a marker of total macrophages (*Cd68*), pro-inflammatory (*Cd38*), or pro-resolving (*Egr2*) macrophages were analyzed. There was a 124.6- and 4.37-fold increase in *Cd68* expression in hyperglycemic BALBc and *Lepr*^db/db^ mice, respectively, suggesting influx of macrophages into the tumor tissue under hyperglycemic conditions. Additionally, hyperglycemic mice had higher expression levels of both *Cd38* and *Egr2*. Similarly, expression of pro-inflammatory markers was upregulated in tumors from hyperglycemic *Lepr*^db/db^ mice (Fig. 2b). Levels of *Tnf* mRNA were significantly increased compared to a normoglycemic *Dock7^m^+/Lepr*^db^ heterozygote control group (*Lepr^Dock/db^*). Macrophage pro-inflammatory marker *Cd38* was increased, but not statistically significant, in *Lepr*^db/db^, which may explain why both *Il6* and *CCl2* expression were upregulated in hyperglycemic mice, but not statistically significant. In contrast, WT mice on a Western diet tended to decrease the expression of each pro-inflammatory marker (Fig. 2c). Expression of *Cd38* was significantly decreased (*P*=.04), which may explain the overall decreased inflammatory expression in these tumors.

**Figure 2.**
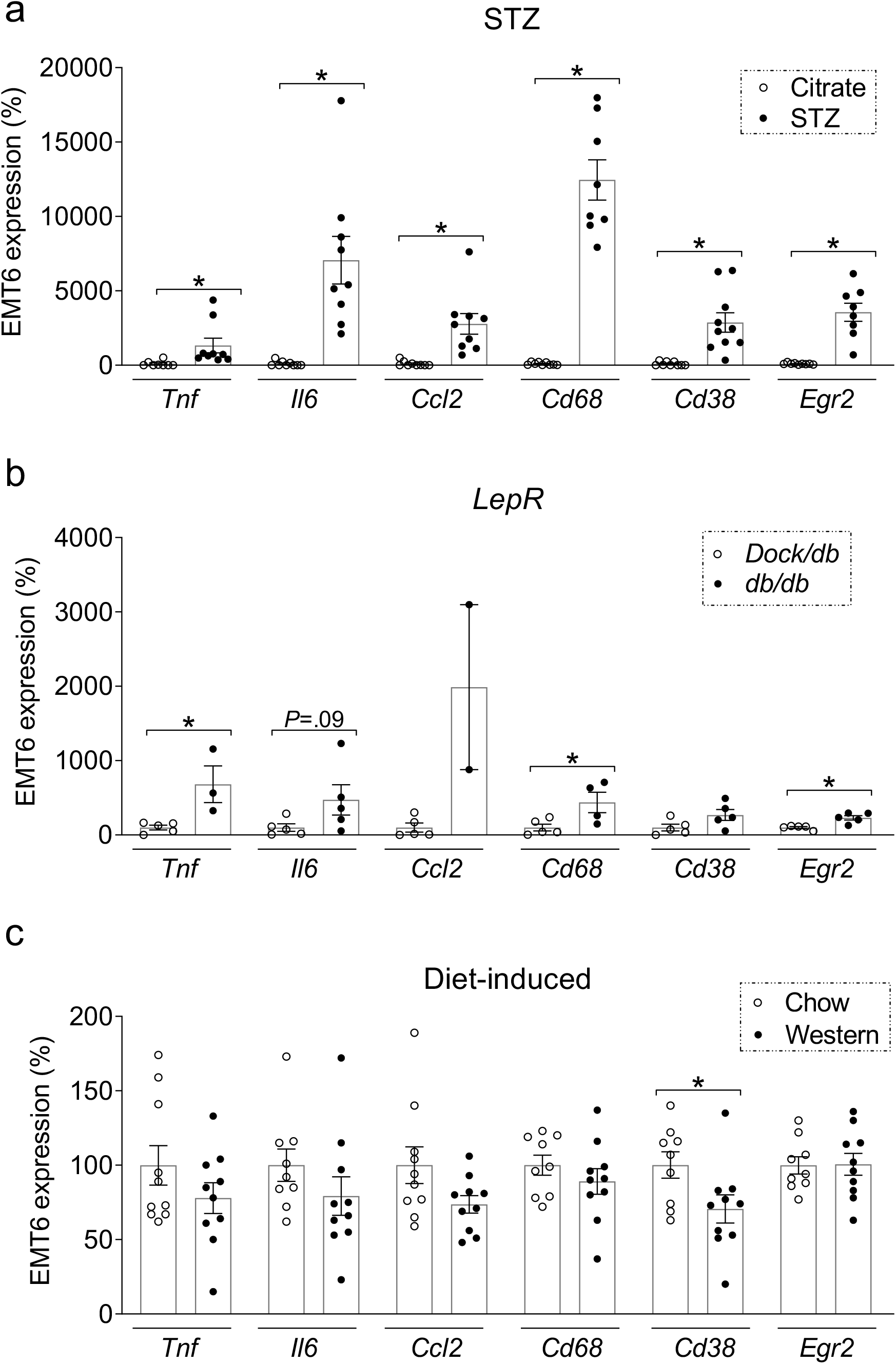
Increased expression of inflammatory markers in EMT6 BC tumors from hyperglycemic mice. RNA from whole tumors were extracted and assayed for pro-inflammatory markers (*Il6*, *Ccl2*, *Tnf*) and macrophage markers (*Cd68*, *Cd38*, and *Egr2*) in: a. BALB/c mice (n=10/group), b. *Lepr* mice (n=5/group), c. WT mice on long term Western diet (n=10/group). Data is normalized to STX5. Bars represent the mean ± SEM. * *P*<.05

**Figure 3.**
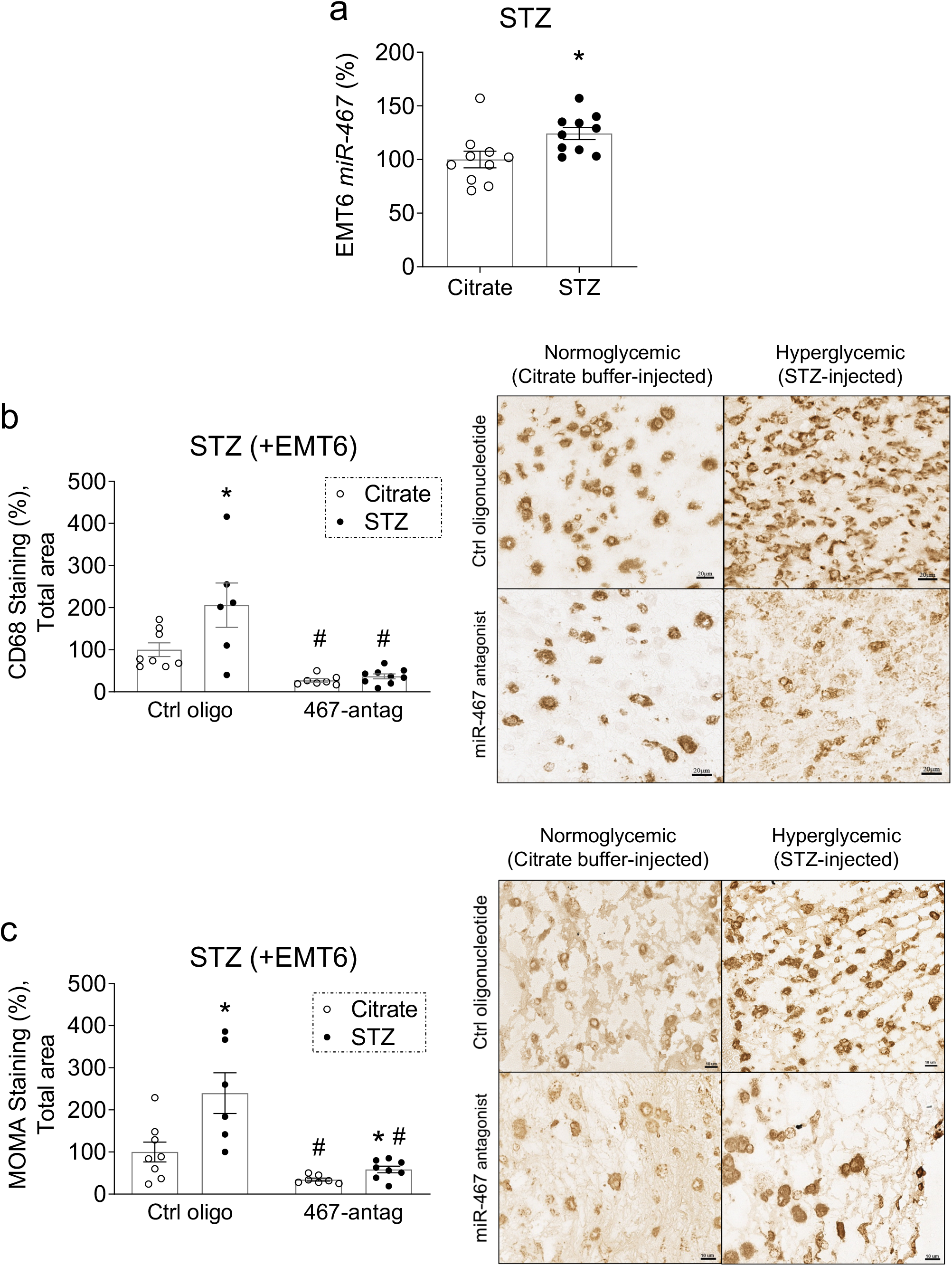
Inhibition of miR-467 with antagonist prevents hyperglycemia-induced macrophage accumulation. a. RNA from whole tumors was extracted and miR-467 was assessed in BALBc mice (n=10/group). b. Sections of tumors harvested from hyperglycemic mice (STZ-induced) and normoglycemic mice that received systemic injections of miR-467 antagonist (or a control oligonucleotide) were stained with anti-CD68 antibody. c. IHC using anti-MOMA-2 antibody to stain for macrophages. The % of stained area was quantified per total section area and multiplied by tumor weight (n=10/group). Scale bars=10μM. Bars represent the mean ± SEM. * *P*<.05 comparing normoglycemic groups, # *P*<.05 compared to ctrl oligo.

### Inhibition of miR-467 prevents macrophage accumulation in BCs from hyperglycemic mice

To understand how miR-467 affects macrophage accumulation in BC, a miR-467 antagonist was used in BALB/c mice injected with STZ, and sections of EMT6 tumors were stained with antibodies against markers of macrophages, anti-CD68 and MOMA-2. Tumors from hyperglycemic mice had a 24.3% increase in miR-467 levels (Fig.3a, *P*=.01) and the macrophage infiltration was also increased (Fig. 3b,c). In the group that received injections of the control oligonucleotide, we detected a 2.06-fold increase in CD68 macrophage staining in BC sections (Fig. 3b, *P*=.03) and a 2.4-fold increase in staining with the MOMA-2 antibody (Fig. 3c, *P*=.02) in hyperglycemic mice (STZ-treated). In normoglycemic mice, CD68-positive staining decreased 3.65-fold in response to injections of the miR-467 antagonist (Fig. 3b, *P*=.001), and MOMA-2 staining decreased 2.89-fold (Fig. 3c, *P*=.02). Inhibition of miR-467 with an antagonist, significantly blunted CD68 macrophage staining in hyperglycemic (STZ-injected) mice by 5.56-fold (Fig. 3b, *P*=.02) and also decreased MOMA-2 staining by 4.08-fold (Fig. 3c, *P*=.01).

### Expression of miR-467 in human breast tissue positively correlates with glucose levels

In collaboration with the Cleveland Clinic biorepository, we obtained normal and malignant breast tissue from chronically hyperglycemic (HbA1c > 7) or normoglycemic (HbA1c < 6) patients. These specimens were assessed for miR-467 expression to determine whether the miR-467-dependent pathway is present in humans. There was a 3.7-fold increase in miR-467 expression in normal breast tissue, comparing hyperglycemic and normoglycemic groups (Fig. 4a, *P*=.02). In normoglycemic patients, there was no difference in miR-467 levels in malignant vs normal breast tissue. In hyperglycemic patients, miR-467 expression was dramatically increased 258-fold in malignant tissue when compared to normoglycemic patients (*P*<.001) and increased 56-fold compared to normal tissue (*P*=.008). Additionally, there was a positive correlation between increasing glucose levels and miR-467 expression in normal breast tissue (Fig. 4b, r^2^=0.6109).

**Figure 4.**
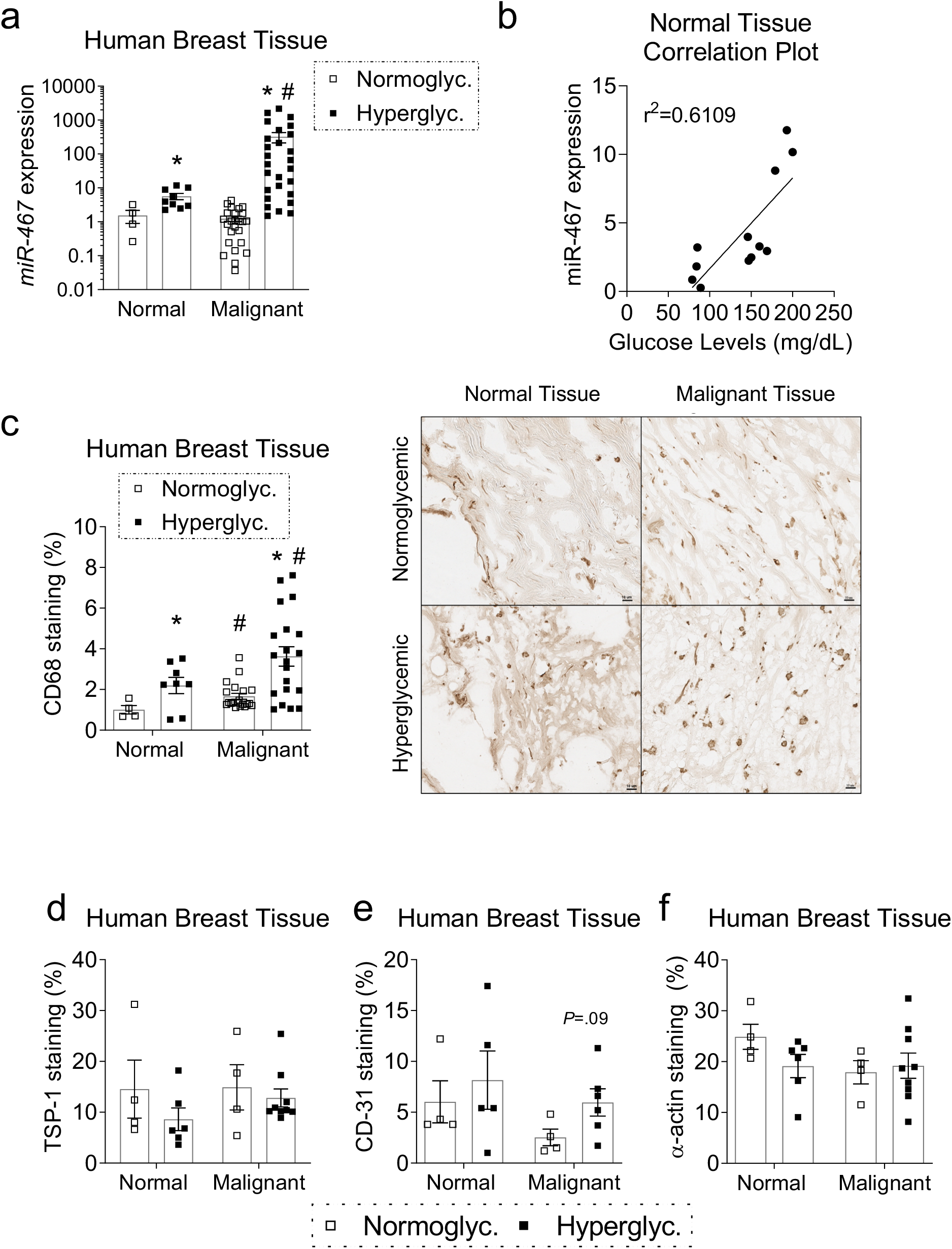
Expression of miR-467 is increased in EMT6 tumors from chronically hyperglycemic patients. a. miR-467 expression was assessed in human breast tissue acquired from the CCF Biorepository. Normal tissue is non-malignant adjacent tissue. Data is log transformed. b. Correlation plot between miR-467 expression and glucose levels in normal, adjacent tissue. c. Sections from human breast tissue were stained for macrophage marker (CD68) and the % of stained area was quantified per total section area and multiplied by tumor weight. Scale bars=10μM. Sections from human breast tissue were stained for TSP-1 (d), and two angiogenesis markers CD31 (e) and α-actin (f). The % of stained area was quantified per total section area. Bars represent the mean ± SEM. * *P*<.05 comparing normoglycemic groups, # *P*<.05 comparing normal tissue.

### Increased macrophage accumulation in malignant breast tissue from hyperglycemic patients

To understand how macrophage accumulation in BC tumors is changed by hyperglycemia, sections of normal and malignant breast tissue samples from patients were stained for a marker of macrophages, CD68 (Fig. 4c). Comparing breast tissue from hyperglycemic patients, macrophage staining increased in both the normal tissue (2.18-fold, *P*=.04) and malignant tissue (2.17-fold, *P*=.002). Macrophage staining was significantly increased in malignant tissue as compared to normal tissue in both normoglycemic (1.65-fold, *P*=.02) and hyperglycemic patients (1.64-fold, *P*=.04).

### Increased angiogenesis and decreased TSP-1 in human diabetic BC samples

To determine whether the negative regulation of TSP-1 by miR-467 is present within the human specimens, breast tissue specimens from patients with HbA1>7 were assessed for TSP-1 protein levels (by an anti-TSP-1 antibody) and levels of the angiogenesis markers (anti-CD31 and anti-α-actin antibodies) (Fig. 4d-f). As expected, staining of TSP-1 tended to decrease along with an increase in CD31 staining, a marker of endothelial cells, in hyperglycemic patients compared to the normoglycemic group. α-actin, a marker of smooth muscle cells and maturation of the vessels, was either decreased in normal breast tissue or not changed in tumor tissue of hyperglycemic patients (Fig. 4f), suggesting growth of less mature blood vessels in hyperglycemic tissues. In normoglycemic patients, α-actin staining was lower in tumors as compared to the normal tissue, consistent with the less mature leaky blood vessels expected in cancer tissue.

### Increased expression of pro-inflammatory markers and markers of macrophages in BC from hyperglycemic patients

Similarly to mouse tumors, expression of macrophage and pro-inflammatory markers were analyzed in tumors from hyperglycemic patients (Fig. 5). Expression of pro-inflammatory markers *IL6* and *CCL2* were increased in tumors compared to the normal breast tissue (Fig. 5a,b). *IL6* levels were further increased 5-fold in malignant specimens from hyperglycemic patients compared to the normoglycemic group (Fig. 5a, *P*=.03). To further understand the type of the increased macrophage population present in these tumors, markers of pro-inflammatory (*CD38,* Fig. 5c) and pro-resolving (*EGR2,* Fig. 5d) macrophages were measured. Both populations were increased in tumors.

**Figure 5.**
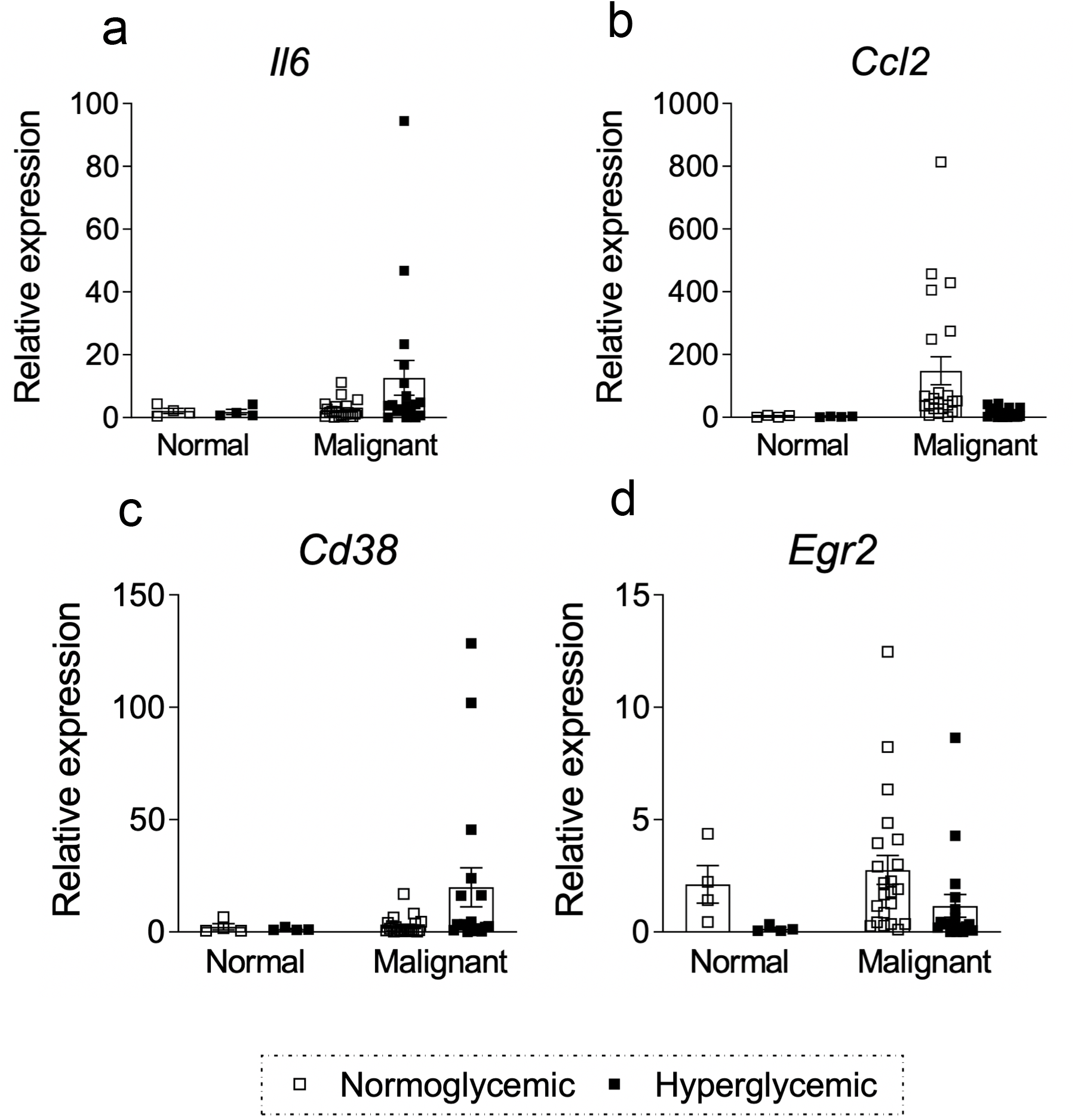
Expression of inflammatory markers in BC tumors from hyperglycemic patients. RNA from whole tumors (acquired from the CCF Biorepository) were extracted and assayed for pro-inflammatory markers (*Il6* and *Ccl2*) and macrophage markers (*Cd68, Cd38*, and *Egr2*) in chronically hyperglycemic or normoglycemic patients. Normal tissue is non-malignant adjacent tissue. Data is normalized to 5S. Bars represent the mean ± SEM. * *P*<.05

However, there was a 7.5-fold increase in *CD38* expression in specimens of tumors from hyperglycemic patients as compared to the specimens of tumors from normoglycemic patients (Fig. 5c, *P*=.04), but the levels of pro-resolving (*EGR2)* macrophages marker were decreased 2.4-fold in hyperglycemic tumors (Fig. 5d, *P*=.006), suggesting that infiltrating TAMs are more pro-inflammatory in human BC tumors of hyperglycemic patients compared to normoglycemic patients.

### Level of circulating plasma miR-467 is higher in hyperglycemic patients

We found that miR-467 is a circulating miRNA: we obtained discarded blood plasma samples from both normoglycemic and hyperglycemic patients and quantified the levels of miR-467. Plasma miR-467 in hyperglycemic patients was increased by 80.5% (Fig. 6a, *P*=.04).

**Figure 6.**
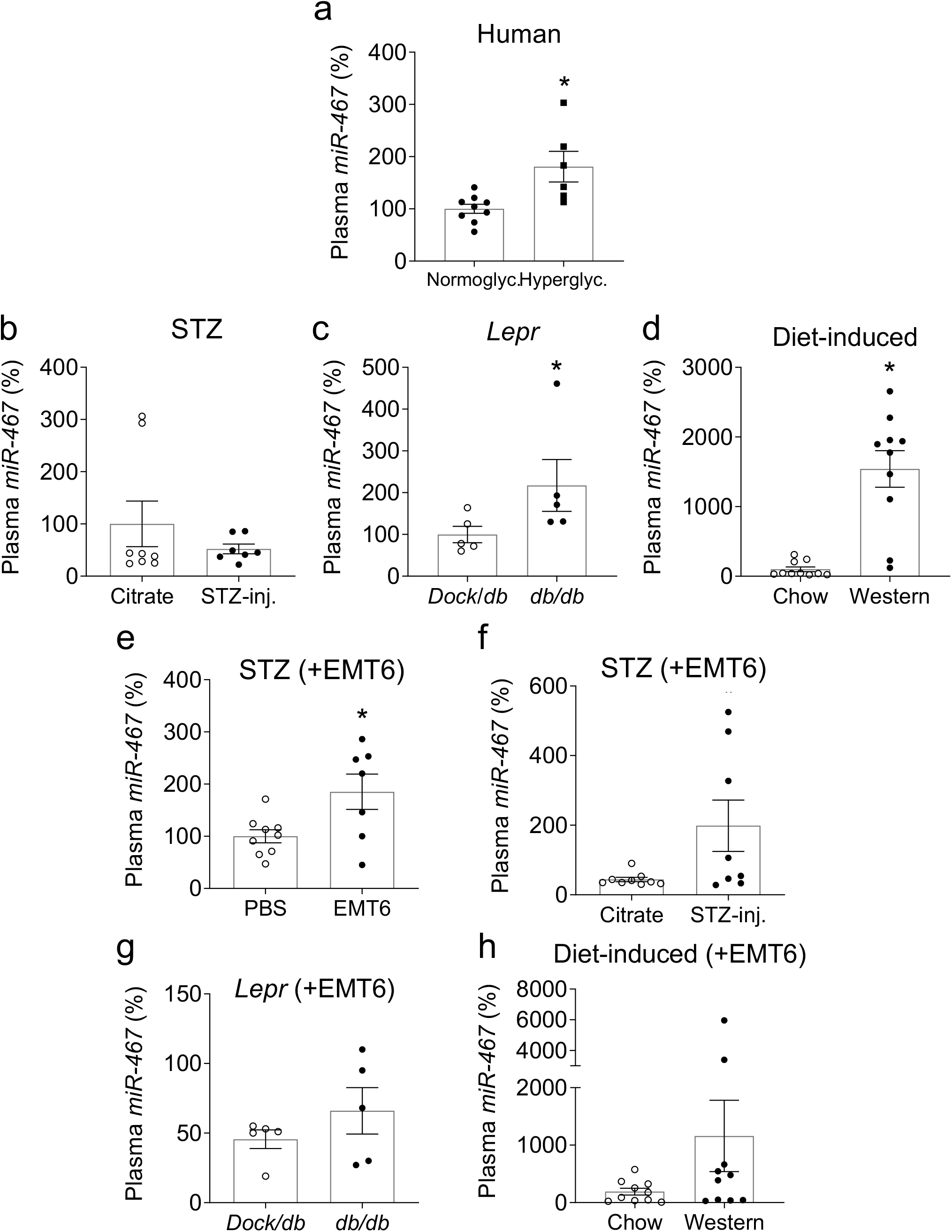
Circulating miR-467 is detected in hyperglycemic patients and mice. a. Plasma from hyperglycemic and normoglycemic patients was acquired from the CCF Biorepository. Plasma miRNA was extracted and miR-467 was quantified by RT-qPCR (n=10/group). b. Plasma from hyperglycemic mice was collected from (STZ-injected) BALBc mice, c. *Lepr* mice, and d. Diet-induced WT mice compared to normoglycemic controls. e. Plasma from hyperglycemic BALBc (all STZ-injected) mice was collected to compare miR-467 in mice injected with BC cells (EMT6) compared to controls. Plasma was collected from mice that were injected with EMT6 cells to determine if circulating miR-467 expression was changed by the presence of the tumor in f. BALBc mice, g. *Lepr* mice, and h. Diet-induced WT mice. Bars represent the mean ± SEM. * *P*<.05 comparing normoglycemic controls.

### Level of circulating plasma miR-467 is higher in hyperglycemic mice

We measured the levels of miR-467 in blood plasma from hyperglycemic mouse models. We did not detect increased levels of miR-467 in STZ-treated BALB/c mice (Fig. 6b), but plasma from *Lepr*^db/db^ mice had a 2.17-fold increase in plasma miR-467 as compared to normoglycemic *Lepr^Dock/db^* heterozygote controls (Fig. 6c, *P*=.03). In a mouse model of diet-induced obesity and chronic hyperglycemia, mice on a Western diet had a 15-fold increase in plasma miR-467 compared to chow-fed mice (Fig. 6d, *P*<.001).

### Level of circulating plasma miR-467 is higher in mice with EMT6 BC tumors

Dramatically increased levels of miR-467 in tumor of hyperglycemic patients and the finding of circulating miR-467 suggested that the levels of circulating miR-467 may be elevated in the presence of a tumor and that miR-467 may clinically useful as a BC biomarker.

We measured the levels of miR-467 in blood plasma samples from hyperglycemic mouse models injected with EMT6 BC cells. In mice injected with STZ to induce hyperglycemia, mice with EMT6 tumors had an 85% increase in plasma miR-467 compared to mice without cancer (Fig. 6e, *P*=.047). We did not detect increased levels of plasma miR-467 in *Lepr^db/db^* and mice on Western diet (data not shown). Because mouse xenografts do not fully imitate the human BC growth, we further explored the changes in miR-467 levels in blood cells and bone marrow (BM) as described below, in order to detect any early signs of miR-467-dependent response in mouse models.

Additionally, the presence of EMT6 BC resulted in increased plasma miR-467 levels in all three hyperglycemia models in response to elevated blood glucose levels: e.g., there was a 4.5-fold increase in plasma miR-467 in STZ-injected hyperglycemic mice with tumors compared to the normoglycemic controls with tumors (Fig. 6f, *P*=.054). In *Lepr*^db/db^ mice with tumors, there was a 46.0% increase in plasma miR-467 compared to normoglycemic mice with tumors, but it did not reach statistical significance (Fig. 6g). In WT mice with tumors on the Western diet, plasma miR-467 had a 6-fold increase compared to chow-fed mice with tumors, although it did not reach statistical significance (Fig. 6h).

### miR-467 levels in whole blood, blood cells, and bone marrow of mice with BC tumors

To further understand the source of circulating miR-467 in response to BC, we analyzed fractions of whole blood for miR-467 in WT mice on a chow or Western diet. This model of hyperglycemia did not have increased levels of miR-467 in plasma, but we expected to find early changes in response to EMT6 tumors. miR-467 levels in whole blood did not change (Fig. 7a). miR-467 levels in red blood cells (RBC) and white blood cells (WBC) fractions were also unchanged (Fig. 7b,c). Lack of changes in miR-467 levels in blood in response to tumor prompted us to look at the effects in bone marrow (BM).

**Figure 7.**
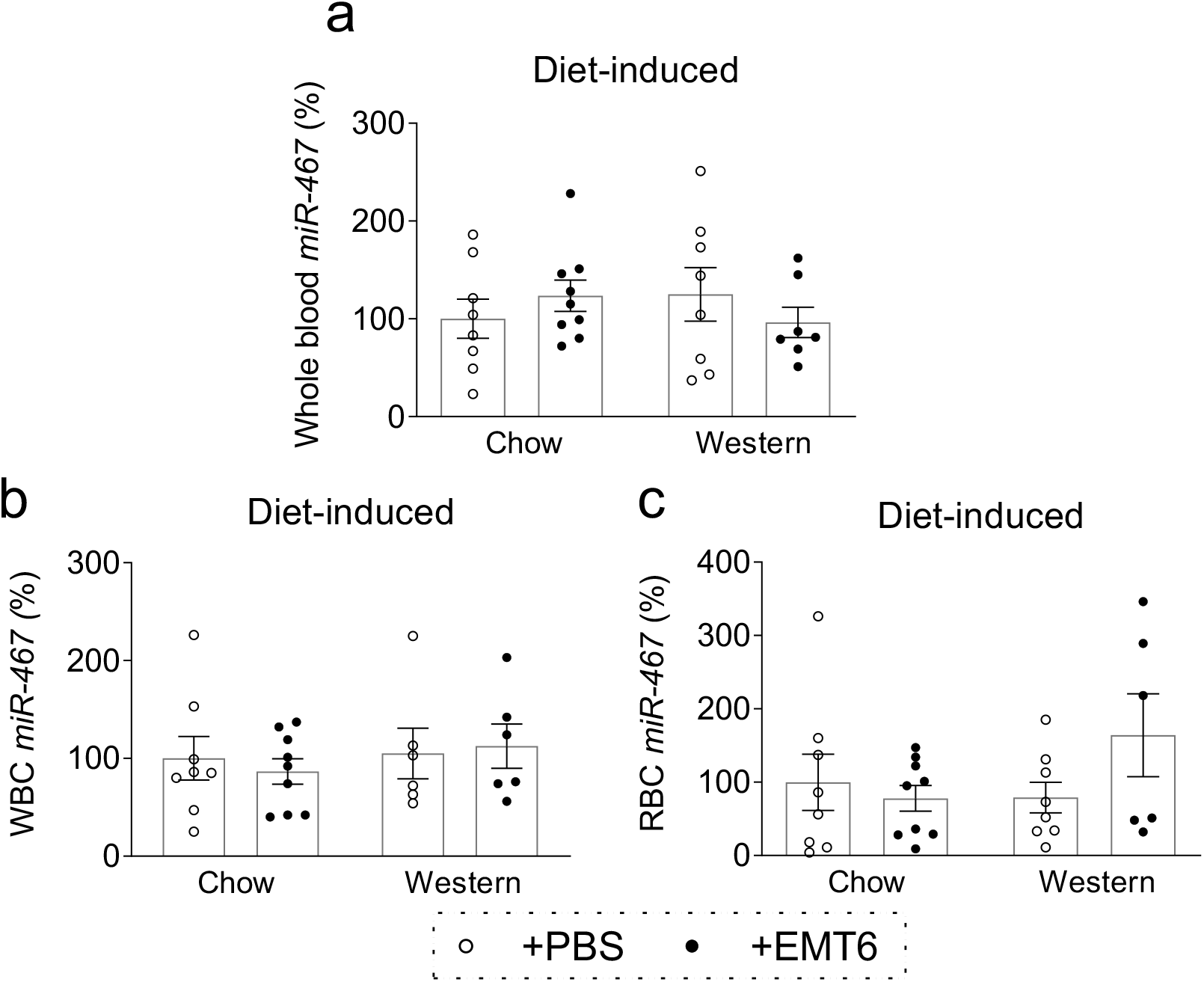
miR-467 expression is unchanged in whole blood and cell fractions in WT mice. a. miR-467 expression was analyzed in whole blood from WT mice on a long-term Western diet and injected with EMT6 BC cells. b. RBC miR-467 expression was analyzed from whole blood that was spun down to remove plasma and the “buffy coat”. c. WBC miR-expression was analyzed from whole blood that was spun down to remove plasma and the buffy coat was extracted to separate out WBC. Bars represent the mean ± SEM.

### Increased miR-467 expression in bone marrow (BM) from hyperglycemic mice

BM analysis demonstrated that, in hyperglycemic mice (STZ-injected), there was a 153% increase in miR-467 levels as compared to normoglycemic mice (Fig. 8a, *P*=.004). A similar effect was observed in WT mice on Western diet: miR-467 was increased by 74.3% in BM (Fig. 8b, *P*<.001). In a genetic model of diabetes, *Lepr* mice, there was no significant differences between the hyperglycemic (*Lepr*^db/db^) group compared to the normoglycemic heterozygous control (*Dock/db*) (data not shown). In mice with EMT6 tumors, we detected a significant increase in miR-467 levels in response to hyperglycemia (Figs. 8c-d). In BM from STZ-treated hyperglycemic mice with tumors, there was a 2.63-fold increase in miR-467 expression compared to BM from normoglycemic mice with tumors (Fig. 8c, *P*<.001). A similar effect was observed in Western diet-fed mice: in mice with tumors, miR-467 was increased by 40.0% in hyperglycemic mice as compared to normoglycemic mice on chow (Fig. 8d, *P*<.001).

**Figure 8.**
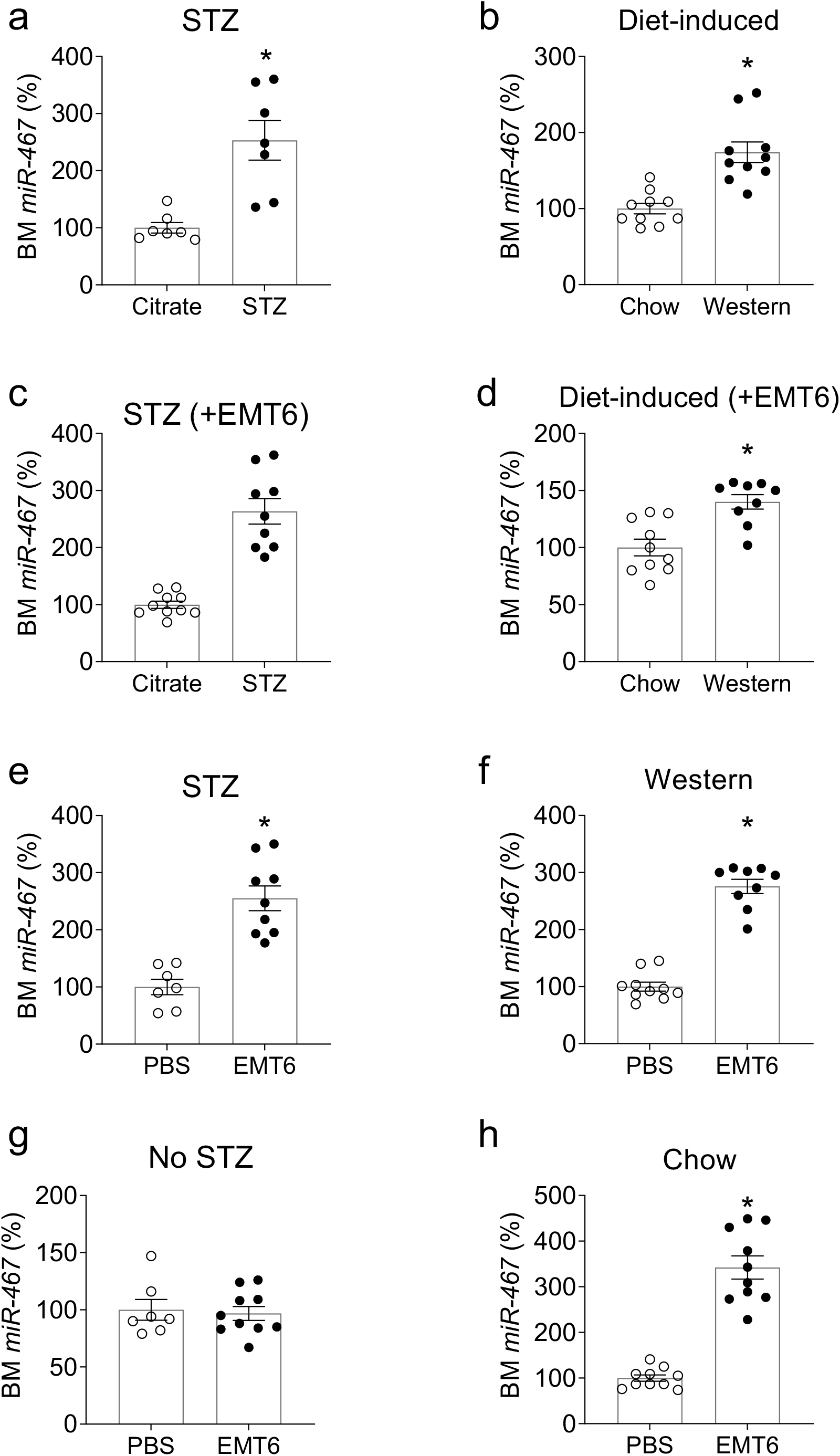
miR-467 expression is increased in whole BM from hyperglycemic mice injected with EMT6 cells. a. Whole BM from hyperglycemic mice was collected from BALBc mice (STZ-injected) and b. Diet-induced WT mice on a long term Western diet to compare miR-467 expression compared to normoglycemic controls. c. Whole BM from mice injected with EMT6 BC cells was analyzed for miR-467 expression in BALBc mice and d. WT C57BL6 mice on a long term Western diet. e. miR-467 expression in BM from hyperglycemic (STZ-injected) BALBc mice or f. WT mice on a long term Western diet was compared in mice with or without EMT6 BC tumors g. miR-467 expression in BM from normoglycemic BALBc mice or h. Chow-fed mice with or without EMT6 BC tumors. Bars represent the mean ± SEM. * *P*<.05 comparing normoglycemic controls.

### Increased miR-467 expression in bone marrow from mice with EMT6 BC cells

The levels of miR-467 in BM were increased in presence of EMT6 tumor as compared to BM from mice without tumors: in hyperglycemic STZ-treated mice with tumors, levels were increased 2.5-fold as compared to BM from hyperglycemia mice without tumors (Fig. 8e, *P*<.001); in mice on Western diet with tumors, the levels of miR-467 were increased 2.7-fold as compared to mice on Western diet without tumors (Fig. 8f, *P*<.001). There was no difference in miR-467 BM expression in normoglycemic BALBc mice with or without tumors (Fig. 8g), but in mice fed a Western diet, presence of BC increased miR-467 expression in BM by 3.4-fold (Fig. 8h, *P*<.001).

## DISCUSSION

Hyperglycemia (HG) is associated with higher incidence of breast cancer (BC). However, patients with metabolic disorders are uniquely understudied and are often excluded from clinical trials and studies. They represent a large fraction of all BC patients: in our study, 18% of all patients had diagnosed diabetes or multiple high blood glucose test results – twice higher than the average incidence of diabetes in the US population. In the US, 331,530 patients are diagnosed with BC every year. Thus, about 60,000 of the new BC patients per year are also diabetic. Furthermore, even intermittent post-prandial elevation of glucose levels in non-diabetic cancer patients is associated with poorer prognosis. Lowering the glycemic index or the glycemic load of meals led to better outcomes in cancer patients (28–52). High blood glucose modulates tissue remodeling, angiogenesis, and inflammation – the events that are intimately related and occur simultaneously in cancer progression. These programs are often regulated through the same or overlapping signaling pathways. Our results indicate that miRNA-467-dependent pathway regulates cancer growth, angiogenesis, and infiltration of TAMs in response to HG.

In the last 25 years, since the discovery of miRNAs, it has become clear that these small non-coding RNAs regulate many key steps in physiology and in the development of various pathologies, including BC (53–55). miRNAs have been studied as prospective cancer markers and therapeutic agents (54, 55).

Currently, there is no safe, reliable, and inexpensive method to detect BC tumors, especially recurrence and metastasis. Mammograms are used routinely, but they do not detect some types of cancer, cannot detect metastasis, and are not without a risk to the patients’ health, which limits their use to once a year - once in two years. MRI is too expensive (and unsafe to use routinely) and ultrasound or other imaging methods are not reliable. The current strategy is “wait and see”: oncologists advise to seek medical help if or when symptoms develop. However, by then, the disease has progressed. A test that can be performed routinely to monitor or even suggest recurrence or metastasis would be extremely helpful. Our results suggest that miR-467 may prove to be a useful marker of a BC tumor. Even if the marker and the mechanisms studied in this work are only applicable to the diabetic patients, it will still benefit hundreds of thousands of patients.

In this study, we report that miR-467 accelerates tumor growth by inducing promoting the recruitment of TAMs to drive HG-induced cancer inflammation. Our initial study of miR-467 as a regulator of expression of a potent anti-angiogenic protein thrombospondin-1 (TSP-1) lead to a discovery of pro-angiogenic function of miR-467 in tumors (25, 56) that is mediated by downregulation of TSP-1 (26). Here, we described its pro-inflammatory function that is mediated by the downregulation of TSP-1 as well as other unknown miR-467 targets.

We used three mouse models to study the effects of increased blood glucose levels on mouse BC xenograft growth, miR-467 levels, and inflammation: 1) STZ-induced hyperglycemia that imitates human type 1 insulin-dependent diabetes, 2) genetic model of insulin resistance and type 2 diabetes (*Lepr^db/db^* mouse), and 3) a diet-induced insulin resistant and hyperglycemic model that may be closest to mimicking the human diet-induced metabolic changes observed in the “Western” population. In all three models, hyperglycemia was associated with the accelerated BC tumor growth. As we previously reported, administration of a miR-467 antagonist prevented the effect of hyperglycemia on tumor size (26). In addition to mouse models, diabetic patients’ specimens of BC tumors and normal adjacent tissues were examined.

The expression of miR-467 in tumors was upregulated in HG: we have detected a significant increase in EMT6 tumors of three mouse models of hyperglycemia and in hyperglycemic patients’ normal breast and cancer tissue specimens. Interestingly, the expression of miR-467 was also upregulated in the bone marrow of hyperglycemic mice, suggesting that miR-467 may be involved in regulation of production of the inflammatory blood cells or regulation of their functions.

HG was associated with the increased levels of several pro-inflammatory markers, including several cytokines (*Il6*, *Ccl2*, and *Tnf*) and *Cd68* (a marker of macrophages). Immunohistochemistry with anti-macrophage antibodies (anti-CD68 and MOMA-2) revealed increased numbers of macrophages in tumors of HG mice, and infiltration of tumor-associated macrophages (TAMs) was prevented by administering a miR-467 antagonist. Both the markers of pro-inflammatory macrophages (*Cd38*) and tissue-repair macrophages (*Egr2*) were upregulated in tumors of hyperglycemic mice suggesting that the polarization of the recruited macrophages was not regulated by miR-467. In patients’ tissues, TAM infiltration was higher in both the normal and the cancerous tissues from hyperglycemic patients as compared to specimens from the normoglycemic patients. Furthermore, TAMs were significantly upregulated in tumors vs adjacent non-cancerous “normal” tissues, supporting more inflammation in tumors, and even further increased infiltration of immune cells in hyperglycemic patients. TAMs contribute to the progression of many cancers (31, 38, 47, 48). Patients with cancers that have higher TAM numbers have lower survival rates (36). Clinically, expression of the macrophage marker CD68 is routinely used as a part of the OncotypeDX test that predicts BC aggressiveness and metastasis (57, 58). In a tumor microenvironment, TAMs (together with other leukocytes) secrete a variety of pro-inflammatory factors and promote chronic inflammation and angiogenesis, thus, supporting tumor growth. The inflammation is increased in hyperglycemic patients, and diabetes is associated with a poorer prognosis and a lower disease-free survival of patients with BC (4–6, 41).

miR-467 levels were higher in hyperglycemic patients’ specimens in both the non-cancerous and cancerous tissues. However, in malignant tissues, the effect of hyperglycemia on miR-467 levels was disproportionally higher, increased by three orders of magnitude as compared to either malignant normoglycemic or non-cancerous tissues from either normoglycemic or hyperglycemic patients. This observation lead us to investigate whether miR-467 circulates in blood and whether the presence of a BC tumor increases the levels of circulating blood miR-467. miR-467 was easily detected in human plasma and its levels were upregulated in the plasma of patients with HG. In three mouse models of hyperglycemia (STZ injections, *Lepr^db/db^*, and Western diet), hyperglycemic mice had higher levels of miR-467 in plasma. When the plasma levels of miR-467 were compared in mice with and without EMT6 tumors, the mice with cancer had higher levels of circulating miR-467, and the levels of circulating miR-467 was consistently higher in hyperglycemic mice with tumors compared to normoglycemic mice with tumors. These data suggest that miR-467 may prove to be a marker of BC in diabetic or chronically hyperglycemic patients. Such marker could be used as an efficient and inexpensive test to monitor metastasis and recurrence after treatment of the primary tumor, similar to the marker that exists for prostate cancer patients. miR-467 circulates in blood and may become a valuable BC tumor biomarker that could be used to detect primary tumor or, more importantly and clinically relevant, cancer recurrence and metastatic disease. miR-467 can be easily measured in the blood to indicate primary or to monitor secondary BC tumors.

To understand the source of circulating miR-467 in plasma, we evaluated miR-467 levels in blood cells and bone marrow of mice with tumors. We did not detect changes in circulating blood cells, but changes in miR-467 levels in BM in response to the EMT6 tumor occurred within days, suggesting that circulating miR-467 may reflect the increased miR-467 levels in specific sub-fractions of BM and the degree of infiltration of tumors with immune cells from BM.

Our study has documented the effects of HG on BC growth and inflammation. There is currently very little information on the mechanisms, by which glucose attracts and/or retains TAMs and promotes a pro-inflammatory environment in tumors. Our data suggests that miR-467, upregulated by high glucose, is a regulator of TAMs and cancer inflammation and may prove to be a clinically useful marker of a BC tumor.

## MATERIALS AND METHODS

### Experimental animals and protocols

All animal procedures were performed according to protocols approved by the Institutional Animal Care and Use Committee and in agreement with the National Institutes of Health *Guide for the Care and Use of Laboratory Animals*. Animals were housed in AALAC approved animal facilities of the Cleveland Clinic. Wild type C57BL6, BALB/c, *Lepr*^db/db^*, Dock7^m^+/Lepr*^db^ (heterozygote control, lean and normoglycemic), and *Thbs1^−/−^* mice were purchased from The Jackson Laboratories. WT C57BL6 mice were fed a chow or Western diet (TD.88137, 40-45% kcal from fat, 34% sucrose by weight, Envigo) starting at 4 weeks of age.

Animals were sacrificed at end point by exsanguination under anesthesia with ketamine/xylazine (100 mg/15 mg/kg), and organs were collected. Mice used for cell isolation were euthanized by CO_2_ asphyxiation followed by the cervical dislocation.

### Induction of Diabetes in Mice

WT C57BL6 and BALB/c mice were given intraperitoneal streptozotocin (STZ; in 20 mmol/L citrate buffer, pH 4.6) injection following the procedure proposed by Jackson Laboratories (50 mg/kg for 5 consecutive days). Age matched controls received citrate buffer injections. Blood glucose was measured starting 48 hours after the final STZ injection using the AlphaTrak Blood Glucose monitoring system. Mice with blood glucose >250 mg/dL were selected for the described experiments. In the *Lepr^db/db^* mice and the control *Dock7^m^+/Lepr*^db^ mice, blood glucose levels were measured at the end of the experiment.

### Injection of cancer cells and tumor collection

EMT6 mouse cells were purchased from American Type Culture Collection (ATCC) and cultured according to ATCC directions. Cancer cells were injected as described in our prior reports. On injection day, cultured EMT6 cells were washed twice with sterile PBS, viable cells were counted, and cancer cells were injected into mammary fat pad (1.5 × 10^6^ cells in 100 μL sterile PBS). Tumors were harvested when the largest tumors in hyperglycemic mice reached the maximum allowed size (1.7mm^3^) on day 10-11 post injection. Tumors were weighed, frozen in OCT, or processed immediately to isolate RNA.

### Immunohistochemical staining

Using VECTASTAIN ABC-HRP Kits (Vector Labs), 10 μm sections were stained with antibodies against CD68 (biotinylated clone FA-11, 1:10, AbD Serotec), CD31 (1:100, BD Pharmingen), laminin (1:300, Abcam), α-actin (clone ab5694 1:200, Abcam), or anti-TSP-1 Ab4 (clone 6.1 1:100, Thermo). Secondary antibodies were included in the species-specific kit, followed by ImmPACT DAB peroxidase substrate (Vector Labs).

Slides were scanned using Leica SCN400 or Aperio AT2 at 20X magnification. Positive staining in the images was quantified using Photoshop CS2 (Adobe) and Image Pro Plus (7.0).

### RNA isolation and Real-Time Quantitative RT-PCR

RNA was isolated using Trizol reagent (Invitrogen) and quantified using Nanodrop 2000 (Thermo). To measure miR-467a expression, 1 – 2.5 μg of total RNA was polyadenylated using NCode miRNA First-Strand cDNA Synthesis kit (Invitrogen) or miRNA 1st strand cDNA synthesis kit (Agilent). Real-time qPCR amplification was performed using SYBR GreenER™ qPCR SuperMix Universal (Thermo) or miRNA QPCR Master Mix (Agilent). The miR-467a primer (GTA AGT GCC TAT GTA TATG) was purchased from IDT. To measure expression of inflammatory markers, 1 – 2 μg of total RNA was used to synthesize cDNA using the SuperScript First-Strand cDNA Synthesis System for RT-PCR (Invitrogen). Real-time qPCR was performed using TaqMan primers for *Il6, Ccl2*, *Tnf*, *Cd68*, *Cd38, Egr2* (Thermo) and TaqMan Fast Advanced Master Mix (Thermo).

### Statistical Analysis

Data are expressed as the mean value ± SEM. Statistical analysis was performed with GraphPad Prism 8 Software. Student’s t-test and ANOVA were used to determine the significance of parametric data, and Wilcoxon rank sum test was used for nonparametric data. A *P* value of <.05 was considered statistically significant.

## ACKNOWLEDGEMENTS

This work was supported by the National Institutes of Health awards R01 HL117216 and R01 CA177771 (O. S. A.) and by the American Heart Association award 17PRE33660475 (J. G.).

## Notes

### Competing Interest Statement

The authors have declared no competing interest.

